# RDAforest: identifying environmental drivers of polygenic adaptation

**DOI:** 10.1101/2024.10.21.619525

**Authors:** Mikhail V. Matz, Kristina L. Black

**Affiliations:** Department of Integrative Biology, University of Texas at Austin; the National Center for Atmospheric Research, Boulder, CO, USA

## Abstract

Identifying environmental gradients driving genetic adaptation is one of the major goals of ecological genomics. We present RDAforest, a methodology that leverages the predominantly polygenic nature of adaptation and harnesses the versatility of random forest regression to solve this problem. Instead of computing individual SNP-environment associations, RDAforest seeks to explain the overall genetic covariance structure based on multiple environmental predictors. By relying on random forest instead of linear regression, this method can detect non-linear and non-monotonous dependencies as well as all possible interactions between predictors. It also incorporates a novel procedure to select the best predictor out of several correlated ones, and uses jackknifing to model uncertainty of genetic structure determination. Lastly, our methodology incorporates delineation and plotting of “adaptive neighborhoods” - areas on the landscape that are predicted to harbor differentially adapted individuals. Such maps can be used as a guide for planning conservation and ecological restoration efforts. We demonstrate the use of RDAforest in two simulated scenarios and one real dataset (North American gray wolves).

## Introduction

Landscape genomics seeks to plot genetic adaptation on a spatial map, by associating environmental parameters with allele frequencies across the genome (Balkenhol et al. 2017). Landscape genomics methodologies are still under active development, but thus far most of them relied upon linear associations between allele frequencies at individual SNPs and environmental predictors (Forester et al. 2018; Caye et al. 2019; Duruz et al. 2019; Privé et al. 2020). The major problem with such an approach is that most adaptive responses in nature are polygenic, based on small allele frequency changes across a great number of loci (Barghi, Hermisson, and Schlötterer 2020), and may be undetectable at single-SNP level. SNP-by-SNP analysis is of course necessary when one aims to identify genes and functions putatively involved in adaptation, but this is but one of many applications of landscape genomics. Others include identifying likely environmental drivers of adaptation, predicting where differentially adapted individuals can be found, predicting which locations are at most risk due to changing conditions, planning assisted gene flow, and finding the most suitable location for releasing an individual. Here we show that all these goals can be achieved without computing individual SNP-by-environment correlations. Inability to generate a list of potentially interesting genes is undoubtedly a limitation of our method, but it is a reasonable tradeoff for higher power of other, more practical, applications.

Our first idea is that responses to environmental selection would be reflected in the overall genetic structure, due to selection against migrants between contrasting environments - so called “isolation by environment”, or IBE (I. J. Wang and Bradburd 2014). If migrants outright fail to reproduce in the alternative environment, selection essentially acts as a physical barrier and affects the whole genome, irrespective of the loci actually responsible for adaptation (I. J. Wang and Bradburd 2014). If selection is moderate in intensity and takes a few generations to eliminate maladaptive immigrant alleles, IBE signal would still be visible in a substantial portion of the genome, because there are too few rounds of recombination to eliminate extensive physical linkage to environmentally adaptive loci (I. J. Wang and Bradburd 2014). Only when selection against immigrant alleles is as weak as to take on the order of 10 or more generations would the “classical” pattern emerge, where genetic divergence is observed only at and near the loci directly involved in adaptation (Charlesworth, Nordborg, and Charlesworth 1997). For practical purposes we are primarily interested in strong and moderate selection, since these would be the cases when fitness of relocated individuals is noticeably affected. Still, even in the case of weak selection the adaptation may be detectable on the whole-genome scale if it is highly polygenic, which is expected for physiological traits (Sinnott-Armstrong et al. 2021).

How does IBE affect genetic structure? Imagining all individuals as a multivariate cloud of points defined by genetic distances between them, adaptation to the same environmental conditions would look like a bulge or extension of the cloud, likely aligned with one of its principal components. So, to find such signals, it is possible to analyze the principal components of the genetic distance matrix - genetic principal components (gPCs) - which would simultaneously leverage the polygenic nature of adaptation to boost sensitivity and dodge the need to rely upon individual SNP-environment associations. In support of this idea, a study in maize recently demonstrated that individual SNP-environment association information did not improve prediction of yield under different environmental conditions, compared to just the population structure captured by the first five gPCs (Li et al. 2024). Likewise, in poplar, the prediction of genetic mis-adaptation calculated from randomly chosen SNPs correlated with common garden observations just as well or better than the mis-adaptation predicted from putative SNPs under environmental selection (Fitzpatrick et al. 2021). (We must acknowledge that by historical convention, principal components of a distance matrix are called “principal coordinates”, but since their mathematical meaning is exactly the same (Legendre and Legendre 2012) we will call them genetic principal components, gPCs, throughout this paper).

In asking how the overall covariance structure of a multivariate dataset aligns with external variables, our method is conceptually similar to the redundancy analysis (RDA, (Borcard, Gillet, and Legendre 2011)), which has the same goal. However, the second idea of our method makes it different from the classical RDA: associations with environmental predictors are detected by random forest regressions instead of linear regression. The advantage is twofold. First, random forest can detect non-linear and non-monotonous associations, for example, peak-like dependencies that may be entirely flat from the point of view of linear regression. Second, random forest seamlessly accounts for all possible interactions between predictors without the need to specify them and without loss of power. Of particular use to us is the random forest implementation called gradient forest (Ellis, Smith, and Pitcher 2012), which extends random forest regression to multiple response variables (in our case, multiple gPCs). It also introduces two innovations that greatly help visualization and interpretation of the results. The first one is the concept of turnover curves, which are graphs to visualize the magnitude of change in the response matrix as one moves across the scale of a given predictor. The other innovation is scaling of predictor importances, typically measured as increase in mean square error when the predictor is permuted, in terms of proportion of variation explained by the predictor. For technical details, we refer the reader to the excellent summaries of how RDA (Borcard, Gillet, and Legendre 2011), regression trees (De’Ath 2002), and random forest (Ellis, Smith, and Pitcher 2012) algorithms work.

Existing methodologies have explored either the former or the latter idea, but did not put the two together. There is a landscape genomics method that uses the overall genetic covariance structure to find associations with the environment, the RDA approach by Forester et al (Capblancq and Forester 2021; Forester et al. 2018). However, it still relies on linear regressions to detect associations. On the other hand, gradient forest has been previously used to analyze landscape genomics data (Fitzpatrick and Keller 2015; Bay et al. 2018), but it was applied to individual SNP genotypes instead of gPCs, thus missing the opportunity to capitalize on the overall covariance structure. Also, such an implementation is limited to just a few hundred SNPs, because of limitations of the gradient forest algorithm.

There are two more innovations in our methodology. The first one is the predictor selection procedure, to identify the true driver of genetic structure out of many cross-correlated predictors. It is a common problem with environmental predictors: for example, many standard bioclimatic variables that are routinely used in landscape genomics (e.g., (Bay et al. 2018)) are correlated with each other. Our solution is based on the observation that predictors that don’t actually drive the response but are simply correlated with the true driver diminish in importance at a higher *mtry* setting (Strobl et al. 2008). *mtry* is the number of predictors randomly chosen for tree construction in a specific random forest replicate. Correlated predictors appear important at lower *mtry* because they “stand in” for the true driver that was not chosen in that replicate. At higher *mtry*, the true driver is more often present among the chosen predictors and is then preferred for tree construction, which leads to diminishing importance of “stand-in” predictors. Our procedure reruns the analysis at two *mtry* settings and highlights predictors that do not decrease in importance at higher *mtry*. To the best of our knowledge, this is the first time such a procedure is being implemented in random forest regressions.

The second innovation is the jackknifing procedure when computing gPCs, which models uncertainty of their determination due to finite sample size. In each jackknifing replicate we withhold some proportion (20% by default) of all samples from the analysis, compute gPCs without them, and then project the withheld samples onto the ordination to obtain their scores. This models how different the gPCs would be if we had 20% fewer samples.

## Methods

### Inputs

#### Genetic distances

*RDAforest* operates on a square matrix of genetic distances between individuals. This can be computed from a matrix of genotypes (*G*) in [0,1,2] format (i.e. listing the number of derived or non-reference alleles at the given SNP in a given individual) by computing genotypic correlations using R function *cor()* and converting them to distances by subtracting them from 1. For non-model species with Restriction site Associated DNA (RAD) sequencing or shallow whole-genome sequencing (WGS) data, a robust way to compute genetic distances is provided by ANGSD software (Korneliussen, Albrechtsen, and Nielsen 2014): it is identity-by-state (IBS) calculation based single-read resampling. For each base in the genome that is covered by at least one read in each of the two compared individuals, the procedure randomly samples one read from each individual and records either match or mismatch. The proportion of observed matches is the IBS, and the distance returned by ANGSD is 1-IBS (note that the IBS-distance of an individual against itself or its clone is not 0 but half the individual’s heterozygosity). The method does not require genotype calling and is robust to variation in coverage depth between samples. Another promising method for computing genetic distances from raw low-coverage read mapping data is PCAngsd (Meisner and Albrechtsen 2018), which takes genotype likelihoods from ANGSD output and computes the genetic covariance matrix while adjusting genotype likelihoods based on the emerging population structure. The covariance matrix can then be then converted to distances based on genotypic correlations using *cov2cor()* function in R.

#### Environmental variables

The core *RDAforest* analysis uses a table of environmental variables (*E*) with rows corresponding to each individual in *G* (in the same order) and columns corresponding to environmental data. *E* could contain hundreds of variables, some correlated with each other. While *RDAforest* is designed to deal with correlated predictors to a degree, we recommend pruning the most strongly correlated ones, to leave only one representative variable for a group that shows cross-correlations of *r* = 0.9 or higher.

#### Spatial coordinates of genotyped individuals

This is a table (*S*) with two columns giving spatial locations of sampled individuals (with rows in the same order as *G*). It is needed to check whether the genetic structure follows the isolation-by-distance (IBD) pattern and to remove that trend, if present. Typically the spatial coordinates are longitude and latitude. To avoid distortions across the latitudinal range, we convert them to “great circle coordinates”, which are the first two principal components of the great circle distance matrix that takes the curvature of the Earth into account. Great circle distances are used to investigate the presence of IBD, and great circle coordinates are used to correct for IBD (if it is detected) during ordination construction.

#### Prediction space

This is a large table (*P*) containing latitude and longitude coordinates of points where adaptation is to be predicted and their associated environmental values. To keep things simple, we recommend specifying a grid of points on the map at a desired final resolution of the predictions, which would merge into a continuous layer when plotted with a resolution-size symbol.

### Workflow

#### Exploring and countering isolation-by-distance trend

Genetic structure across distances exceeding per-generation dispersal range tends to follow the isolation-by-distance (IBD) pattern, with closely located individuals being more similar genetically (Wright 1943). The easiest way to visually detect this trend is to plot genetic distances against geographic distances, looking for a sloping cline, such as the one on Fig. 3B (or multiple clines, if there are additional drivers of genetic structure besides IBD). To formally test the significance of association between genetic and geographic distances one can use a Procrustes test (function *vegan::protest(), (Oksanen et al. 2007)*) or the classical Mantel test. IBD is a nuisance trend for us because it is not driven by adaptation to specific environmental gradients. To remove IBD signal from the data, we use the fact that on a uniform landscape IBD would produce a match between genetic distances and geographic distances, so that the genetic PCA would look like a geographic map of sampled individuals (House and Hahn 2018). Humans are a good example: their genetic PCA looks very much like a geographic map (Novembre et al. 2008; C. Wang, Zöllner, and Rosenberg 2012). Therefore, to suppress the IBD trend it is sufficient to linearly regress the two spatial coordinates out of genetic data, i.e. construct a partial genetic ordination conditional on the geographic coordinates. This approach is quite conservative since some meaningful environmental gradients may be associated with longitude and especially latitude; regressing spatial coordinates out of the data would de-power our ability to detect the influence of these gradients. Yet, we feel that this should be a default way of dealing with IBD - but only if it is actually evident in the plot of genetic vs geographic distances and in the Procrustes or Mantel test.

#### Constructing genetic ordination

We use the function *vegan::capscale()* (Oksanen et al. 2007) to build the ordination, which accommodates the “Condition” argument to specify nuisance variables to be regressed out of the data (by linear regression). These can be spatial coordinates in case we need to account for IBD, designations of physically isolated regions on the landscape (see Discussion), or technical covariates that might affect the ordination, such as sequencing batch or read coverage. It is highly recommended to explore the shape of the ordination for possible artifacts prior to running the RDAforest analysis, and including all the nuisance variables that appear to have an effect into the “Condition” argument.

#### Exploring the full *RDAforest* model

The goal of this step is to evaluate how good a model we can build with all our predictors in *E*, which gPCs are predictable, which variables in *E* are likely to be the most influential, and which gPCs they are most associated with. At this stage we will deliberately use more leading gPCs that are likely informative (up to all gPCs in the ordination). We do not yet perform any resampling to evaluate uncertainty, this will come later. The function *makeGF()* takes an ordination (partial ordination if IBD was regressed out), table of predictors, and runs *gradientForest()* analysis on it (Ellis, Smith, and Pitcher 2012). This function returns a *gradientForest* model object that contains “importance” (predictive power) of each predictor scaled as proportion of variation explained (R-squared) for each gPC. To compute proportions of total variation in the whole dataset explained by each predictor, per-gPC R-squared values must be summed up while weighing them by the proportion of total variation attributable to each gPC, which is what the function *importance_RDAforest()* does. Summing these values over all predictors gives the cumulative proportion of variation explained by the model. The *$result* slot of the *gradientForest* model contains the list of gPCs that are predicted with a non-zero R-squared; only these gPCs will be used in subsequent stages. The *gradientForest* model can also be used to plot turnover curves for each predictor and for each predictable gPC, which is a good way to visualize which gPCs are responsive to which predictor(s) and where in the predictor’s range major gPC transitions happen.

Note that here as well as everywhere in the *RDAforest* pipeline the R-squared values are computed based on cross-validation, using “out-of-bag” samples that were withheld from model building. This is different from the classical linear regression, where the custom is to report goodness-of-fit measures based on the same data that were used for model building. Such measures are always inflated compared to the true predictive power of the model if it was applied to new data. Cross-validation R-squared values may seem low to new users, but they are a more honest reflection of the model’s predictive power.

#### Predictor selection based on *mtry*

*gradientForest* models are constructed using package *extendedForest*, which already handles correlated predictors to a degree: it employs conditional permutation (Strobl et al. 2008) to avoid over-inflating the importance of correlated predictors. Still, the predictive power remains somewhat spread over several correlated predictors, even if only one of them is truly influential. To identify which one of the group of correlated predictors is the most likely true driver (or most closely correlated with the unidentified true driver), RDAforest uses the *mtry* criterion: at higher *mtry*, true predictors rise in importance, while non-influential correlated predictors diminish in importance, compared to the lower *mtry* setting. This was first seen in simulations that led to the development of the conditional permutation algorithm (Strobl et al. 2008) and confirmed in simulations described below. Additional selection criterion is the relative predictor’s importance: we discard predictors whose importance is less than specified fraction (default 0.1) of the importance of the best predictor. Here, we use *gradientForest*-style importance (R-squared for the predictor averaged over all predictable gPCs) rather than RDAforest-style importance introduced in the previous paragraph (proportion of total variation predicted), because the latter would create bias towards predictors associated with lower-number gPCs.

#### Ordination jackknifing

Predictor selection is where we use ordination jackknifing for the first time. The motivation for this procedure is that with finite sample size the gPC axes are identified with some error, which might affect the apparent importance of predictors. To model the uncertainty of gPCs determination, we run multiple jackknifing replicates where we withhold a certain fraction of the data (default 20%) when constructing gPC axes, thus modeling their uncertainty if we had 20% less samples. The hold-out samples are projected back to the ordination before running the *gradientForest* analysis so we don’t lose power due to smaller sample size. The *mtry*-based selection procedure builds two *gradientForest* models on each replicate, with higher and lower *mtry* setting, and records the importance of each predictor. These results are then summarized as the proportion of replicates in which the importance or a given predictor increases at higher *mtry*.

#### Measuring amount of variation attributable to most important predictors

After the most important predictors are identified, we use them in another series of ordination jackknifing replicates with the function *ordinationJackknife()*, this time recording the importance (proportion of total variation explained) for each passing predictor in each replicate. Boxplot of these values is the end-result of the analysis of the influence of the environment on genetic structure.

#### Predicting adaptation at unsampled locations

We form two types of predictions for the new environmental dataset (*P*): gPC scores predicted by *randomForest()*, and turnover curve values predicted by *gradientForest()*. These predictions are generated by *ordinationJackknife()*, averaging across jackknifing replicates. We recommend running a relatively high number (50 or more) replicates to ensure that predictions are robust across the landscape.

#### Plotting “adaptive neighborhoods”

Earlier papers (Fitzpatrick and Keller 2015; Fitzpatrick et al. 2021; Bay et al. 2018) followed the vignette for the *gradientForest* package (Ellis, Smith, and Pitcher 2012) and plotted turnover curve values as indicators of differential adaptation (or different community composition) on the landscape. However, as we show in simulations below, using turnover curves for this purpose would imply that every transition across a given predictor’s range leads to a new (progressively diverging) state and happens regardless of values of other predictors. Thus, it is appropriate only when changes across the range of each predictor are monotonous, and when there is no interaction between predictors. In contrast, gPC scores predicted by *randomForest()* are much more appropriate to plot since they retain both non-monotony and interactions; however, they may be noisy across space since every point on the landscape is predicted on its own. One way to improve this is by clustering landscape points based on turnover curves, which do integrate over multiple data points to infer boundaries where major transitions happen. Such clusters would robustly group landscape points containing similar adaptation, with a caveat that adaptation within a given cluster can be the same as in some other clusters. To solve this problem, we generate clusters based on turnover curves and then merge clusters that contain similar *randomForest()* predictions.

## Results

### “Thresholds” simulation

In this “toy” simulation (Fig. 1) the distribution of entities (species or alleles) on the landscape follows simple thresholding rules. This is deliberately unrealistic; the primary goal of this simulation is to illustrate how core algorithms of the *RDAforest* work. The environment is a 50 × 50 square grid with two important predictors, *a* and *b*, forming orthogonal gradients. There are additional four predictors correlated with *a* and two predictors correlated with *b*, plus three random predictors (Fig. 1 A, B). The landscape is populated by five “species” that occupy certain regions defined by *a, b*, or their combinations (Fig. 1 C). Note that two of these “species” (“a:b” and “b”) show non-monotonous (peak-like) dependence on the predictors and would not show any linear association with them. The distribution of the “species” on the landscape can be summarized as the PCA map (Fig. 1 D), which shows scores of the first three PCs of the “species” community on the landscape as color mixtures. The higher the color contrast, the more distinct the communities are. To the best of our knowledge, this visualization was introduced in the vignette to the *gradientForest* package, (Ellis, Smith, and Pitcher 2012).

**Figure 1.**
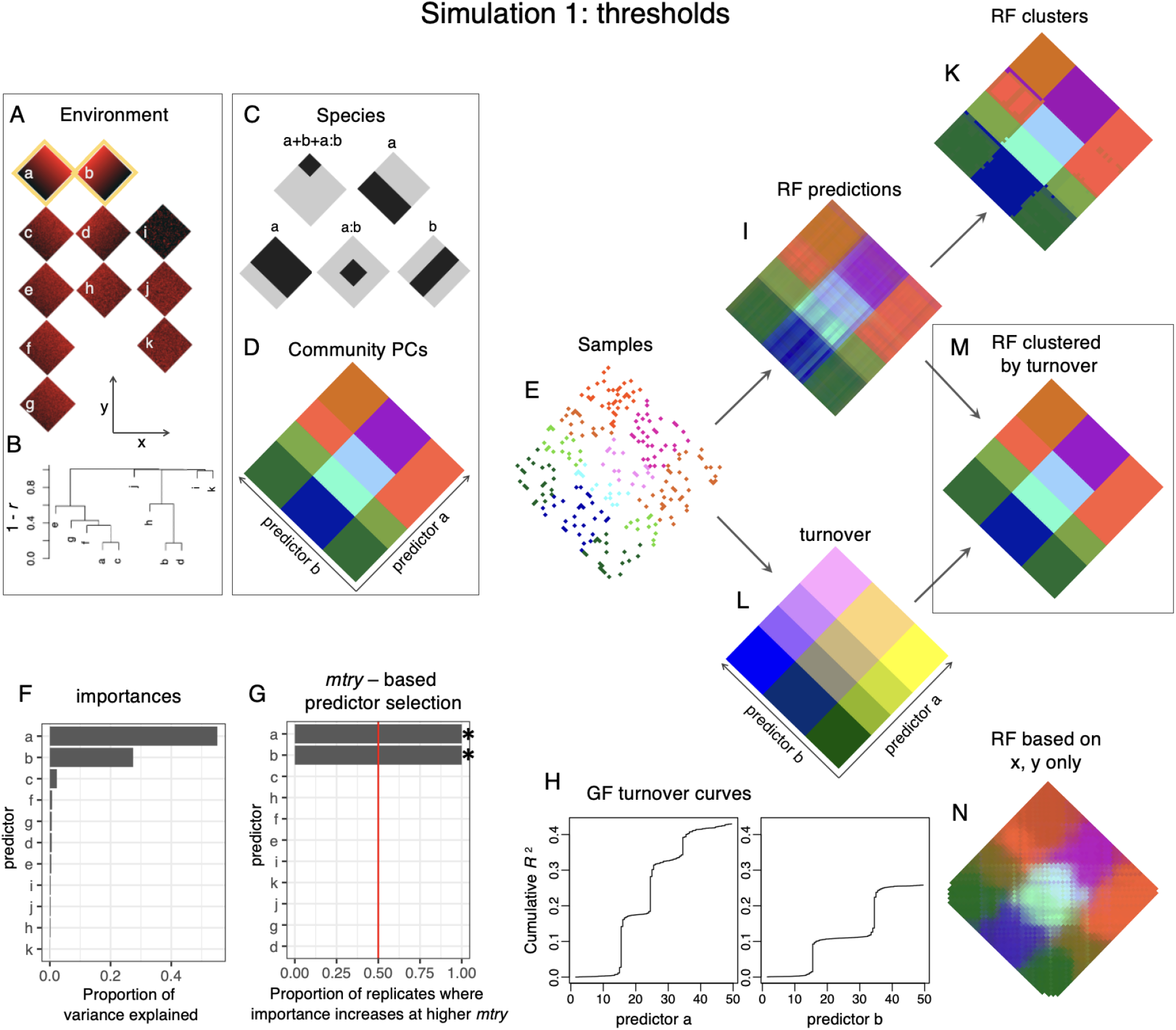
“Thresholds” simulation: a toy example to illustrate how algorithms included in the RDAforest work. A: Environment is a 50 × 50 square with truly influential predictors *a* and *b*. There are four additional predictors (*c, e, f, g*) correlated with *a* and two more (*d, h*) correlated with *b*, plus three random predictors (*i*-*k*). Spatial coordinates are *x* and *y*. B: Hierarchical clustering tree of predictors based on their correlation across the landscape. C: Occurrences of five “species” across the landscape (black - occupied, gray - unoccupied), which in this simulation follows simple threshold rules. Each species occupies regions defined by predictors *a* and *b*. D: PCA map of “communities”. Colors reflect the scores along the first three PCs of the community composition, based on species presences-absences. E: Samples (10% of total) for model building. F: Importances of predictors inferred by a simple *gradientForest* model (no jackknifing). G: Result of *mtry*-based selection procedure, based on 50 jackknifing replicates. Asterisks mark the passing predictors. H: Turnover curves for the passing predictors; steps are the boundaries where the community transitions to a different state. I: PCA map of gPCs scores predicted by *randomForest* (RF), for each point separately. K: Clustering of the RF predictions. L: PCA map of turnover-based predictions. Note that while the boundaries between *potentially distinct* communities are reconstructed quite precisely, the values within each such partition do not reflect their actual differences (compare the color distribution to panel D). M: Clustering of turnover curve predictions followed by merging the clusters based on RF predictions that they contain. N: *randomForest* predictions of the community based exclusively on spatial coordinates (*x* and *y*, panel A), using twice denser sampling than shown on panel E.

The input for *RDAforest* analysis is a random selection of 10% of the landscape points (Fig. 1 E). The *gradientForest* model indicated that predictors *a* and *b* are the most important (in terms of proportion of total variance explained), with *b* less so than *a* (Fig. 1 F). This difference in importance makes sense since there are four transitions along the range of *a* versus three transitions along the range of *b*; moreover, all four regions along *a* are highly distinct while the first and third regions along *b* are similar (compare colors on Fig. 1 D). The *mtry*-based selection procedure robustly identified *a* and *b* as the only viable predictors (Fig. 1 G). Turnover curves for them (Fig. 1 H) reflect the number of transitions (“steps”) along each predictor’s range, which values of the predictor correspond to transition boundaries (*x*-axis coordinates of “steps”), amount of variation attributable to each transition (height of each “step”), and the proportion of total variance explained (the maximum *y*-axis value reached).

The next stage is to use the model based on *a* and *b* to predict (in this case, recreate) the distribution of distinct communities across the remaining 90% of the landscape. Two types of predictions are formed. *randomForest* (RF) predictions (Fig. 1 I) are a direct attempt to predict gPC scores based on the model, for each point individually. Note that RF does a decent job (compare RF predictions on panel I to the true community picture on panel D) but is somewhat uncertain about the exact gPC values, especially at the boundaries of “species” ranges. Clustering these predictions (Fig. 1 K) improves the situation but some mis-predictions along the boundaries remain. In contrast, predictions based on turnover curves made by *gradientForest* (Fig. 1 L) reconstruct the boundaries perfectly, but in many cases the distinctions between bounded regions are incorrect (compare color contrasts on panel L to panel D). There are two problems here. The first one is that turnover-based predictions make it look like the difference always increases with each new transition, which is not necessarily the case: for example, the second transition along the range of *b* leads to almost the same community as the initial state (Fig. 1 D), but in turnover-based predictions it looks like the difference keeps accumulating (Fig. 1 L). The second problem is that turnover predictions assume that each transition for a given predictor happens across the whole landscape, irrespective of other predictors’ values (i.e., predictors don’t interact). For example, in reality the fourth transition along predictor *a* does not happen unless *b* is higher than 15 (Fig. 1 D), but turnover-based predictions miss that. Still, the fact that boundaries between *potentially distinct* communities are reconstructed quite precisely (unlike RF predictions) makes turnover-based predictions useful: we first cluster the turnover-based predictions to identify all the bounded regions, and then merge them based on RF-based predictions contained within the regions. This two-step procedure leads to the perfect recreation of the true community picture (Fig. 1 M).

Last but not least, this simulation is useful to illustrate that one should not attempt to remove variation that can be explained by spatial coordinates alone (*x* and *y*, Fig. 1 A). The random forest algorithm is versatile enough to reconstruct *any* configuration of patches based on just the two linear coordinates (Fig. 1 N), provided dense enough sampling (we have used 20% of all points to generate Fig. 1 N). Clearly, removing this signal would strongly (and perhaps even completely) de-power the main analysis since it also aims to detect *the same patches* on the landscape.

### “Frequencies” simulation

This simulation is more realistic in that the probability of finding certain species (or allele) depending on the predictor changes smoothly across the predictor’s range, instead of simply flipping between 0 and 1 like in the “thresholds” simulation. The environment (Fig. 2 A, B) remains the same as in the “thresholds” simulation, but the “species” (Fig. 2 C) communities now show smooth transitions instead of sharp boundaries. There is also one “species” that is found everywhere, to avoid having points with no species present. Note that the spatial distribution of true PCA values of the community (Fig. 2 D) is very noisy, reflecting the graininess of the simulation and the stochasticity of species occurrences. Our goal will be to draw the most reasonable community boundaries despite this noise.

**Figure 2.**
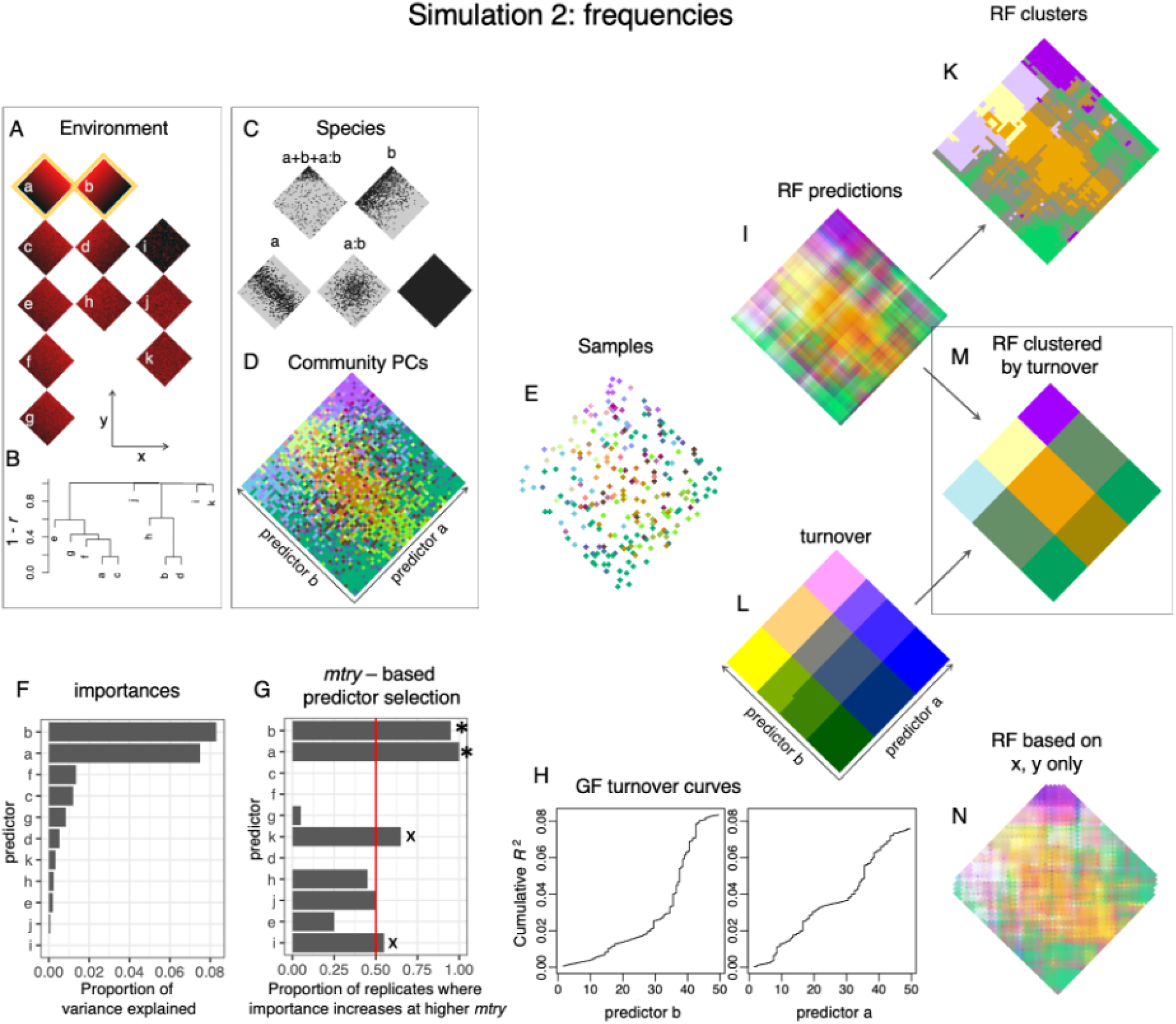
“Frequencies” simulation: a more realistic example. A and B: Environment is the same as in “thresholds” simulation. C: Occurrences of “species” across the landscape (black - occupied, gray - unoccupied). The probability of finding a species changes smoothly across the predictor range. D: PCA map of “communities”. Colors reflect the scores along the first three PCs of the community composition. The noisiness reflects the stochasticity of species’ occurrence. E: Samples (10% of total) for model building. F: Importances of predictors inferred by a simple *gradientForest* model (no jackknifing). G: Result of *mtry*-based selection procedure, based on 50 jackknifing replicates. Asterisks mark the passing predictors, “x” marks predictors that pass the *mtry* criterion but fail the importance criterion (they explain less than 0.1 of the variation explained by the best predictor). H: Turnover curves for passing predictors. Note that steps here are smoother than in “thresholds” simulation, because transitions between communities are gradual. I: PCA map of gPCs scores predicted by *randomForest* (RF), for each point separately. K: Clustering of the RF predictions. L: PCA map of turnover-based predictions. Note that the boundaries between communities are inferred quite reasonably, despite the noise in actual data (panel D). M: Clustering of turnover curve predictions followed by merging the clusters based on RF predictions that they contain. N: *randomForest* predictions of the community based exclusively on spatial coordinates (*x* and *y*, panel A), using twice denser sampling than shown on panel E.

Using 10% of all points (Fig. 2 E) to build an *RDAforest* model, we first see that, once again, *a* and *b* are identified as the most important predictors (Fig. 2 F). However, the proportion of total variance explained by them is notably lower than in the “thresholds” simulation, because of the stochasticity in species occurrences (Fig. 2 C). The *mtry*-based predictor selection procedure identifies *a* and *b* as the only two viable predictors (Fig. 2 G); note that several additional predictors (marked by “x” symbols on Fig. 2 G) pass the *mtry* criterion but are discarded due to low importance relative to the best predictor. Turnover curves (Fig. 2 H) have smoothened steps reflecting the fact that transitions between communities are gradual, not abrupt like in “thresholds” simulation. The RF predictions (Fig. 2 I, K) recapitulate the noisiness of the original community (Fig. 2 D). With clustering the communities are sometimes reasonably well separated (such as blue and purple in the lower right side of the landscape), but more typically the boundaries between them remain undefined (Fig. 2 K). Turnover-based predictions show better promise in terms of drawing community boundaries (Fig. 2 L) although it is often incorrect in predicting differences between communities, just like in the “thresholds” simulation. Clustering based on turnover predictions followed by merging the clusters based on RF predictions generates a map that demarcates regions containing different communities reasonably well (Fig. 2 M). And once again, *randomForest* based on just the spatial coordinates *x* and *y* (Fig. 2 A) is quite good at predicting the whole *a* and *b*-driven community patchiness (Fig. 2 N), so removing all detectable spatial signal from the data prior to the analysis would not be a good idea.

### Real data analysis (North American wolves)

This dataset comes from (Schweizer et al. 2016) and has been used multiple times to illustrate the power of landscape genomics methods (Capblancq and Forester 2021; Smith et al. 2024). For *G*, there are 94 wolves sampled across Canada and Alaska genotyped for 29,437 SNPs. For environmental predictors for each wolf (*E*), we have a standard set of bioclimatic variables from https://www.worldclim.org/data/bioclim.html, plus tree cover data for the approximate year or sampling (2010) from http://www.cec.org/north-american-environmental-atlas/land-cover-2010-modis-250m/. Each wolf has been assigned an “ecotype” in the original paper (Schweizer et al. 2016); we expect that our analysis will recover those ecotypes as differentially adapted groups. For *P* (prediction space) we have a raster of points covering the whole Canada and USA at one-third of a latitudinal degree resolution (approximately 37 km) and associated with the same variables as in *E*. These data and the Rmarkdown script for the analysis described below are available at https://github.com/z0on/RDA-forest, the Rmarkdown output can be viewed at https://rpubs.com/cmonstr/1268717.

The first step is exploration of genetic structure and isolation-by-distance (IBD). Wolves appear to be highly genetically differentiated across the landscape, apparently with respect to the assigned ecotype (Fig. 3 A). Plotting genetic distances against geographic (great circle) distances revealed a clear sloping cline (Fig. 3 B), signifying prominent IBD; which was also supported by the Procrustes test of association between scores of genetic ordination (Fig. 3 A) and geographic coordinates (function *vegan::protest*, p < 0.001). To remove the IBD trend from the data, we formed a partial ordination conditional on the great circle coordinates, which are the first two principal coordinates of the great circle distance matrix (Fig. 3 C). Notably, Atlantic Forest wolves became quite similar to West forest and Boreal forest wolves, and the Arctic ecotype became clearly different from the High Arctic ecotype. The gPCs from this partial ordination are used in the subsequent steps.

**Figure 3.**
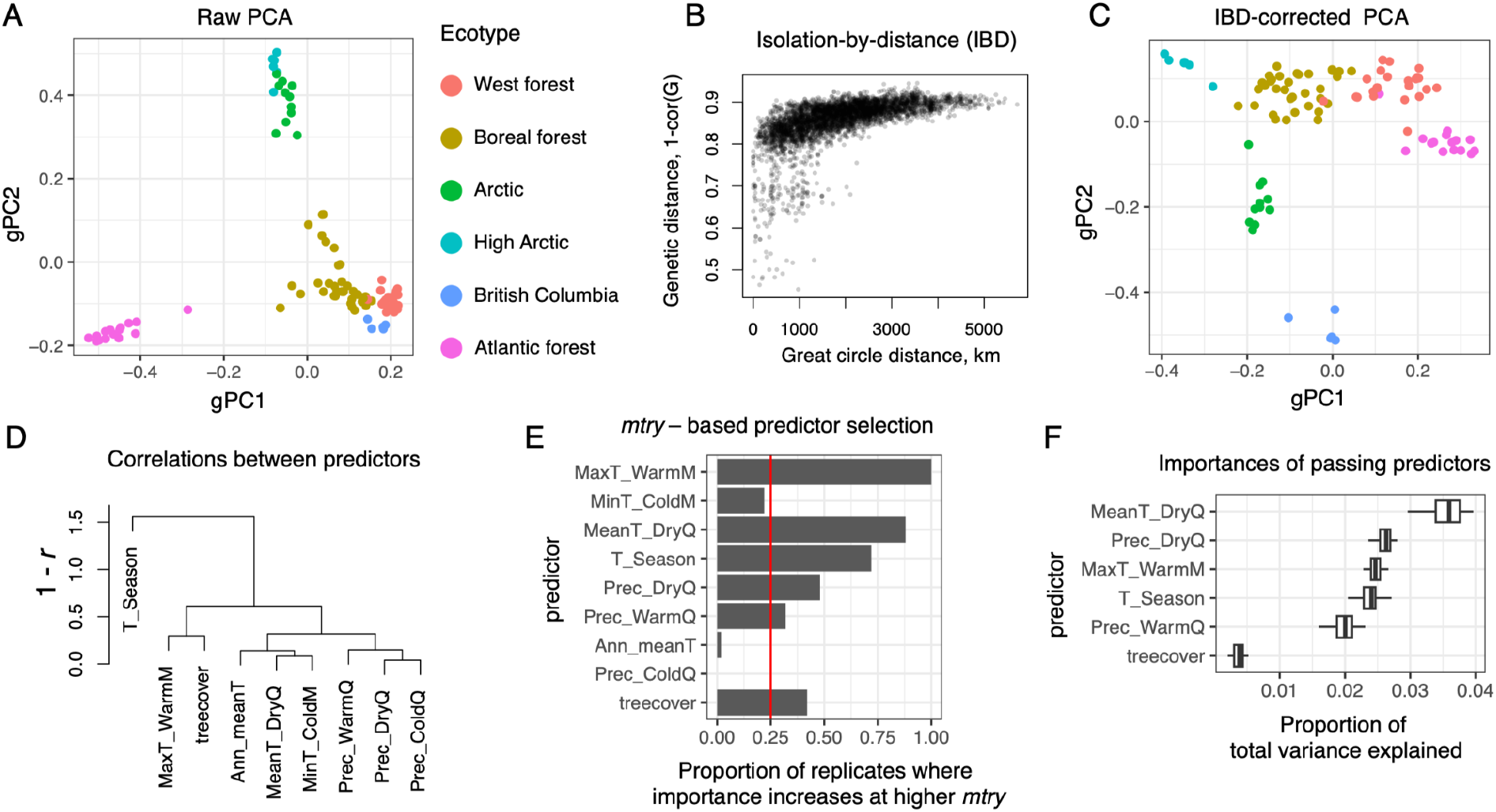
Identifying factors driving genetic adaptation in North American wolves. A: raw PCA of genetic structure, based on genotype correlations. B: Visual examination of the isolation-by-distance (IBD) trend: genetic distances clearly increase with geographic (great circle) distances. C: Partial genetic PCA conditional on the “great circle coordinates”, to correct for IBD. Scores from this PCA become the input for further analysis. D: Correlations between bioclimatic variables and tree cover used as potential predictors. E: Predictor selection based on *mtry*, with somewhat relaxed passing criteria - requiring the predictor to show increased importance at higher *mtry* in only 0.25 of all replicates; red line. F: Boxplots of proportion of variation explained by the passing predictors, based on 50 *ordinationJaccknife* replicates.

The next preparatory stage involves exploration and pruning (if necessary) of predictor variables. We plotted a hierarchical clustering tree of our predictors, based on their correlation, and observed that many bioclimatic variables were strongly correlated with each other. For this analysis, we have removed all but one representative variable out of groups forming clades below 0.1 in the hierarchical clustering tree (i.e., groups of variables correlated with each other with *r* > 0.9, Fig. 3 D).

Next, we perform exploratory modeling. We fit a non-jackknifed gradient forest model with all predictors to 40 leading gPCs, which is deliberately more than we expect to be predictable. The *$result* slot of the model lists gPCs that are actually predictable (i.e. show non-zero cross-validation R-squared). Only these gPCs will be used in subsequent modeling. We then fitted another gradient forest model to the most highly predictable gPCs (the three leading ones) and plotted turnover curves for each predictor:gPC combination (Fig. 4). These plots help visually assess which predictor is associated with which gPC, and where across the predictor’s range major gPC transitions happen. In addition to being a good “sanity check” for the dataset, such visualization greatly improves the interpretability of the model.

**Figure 4.**
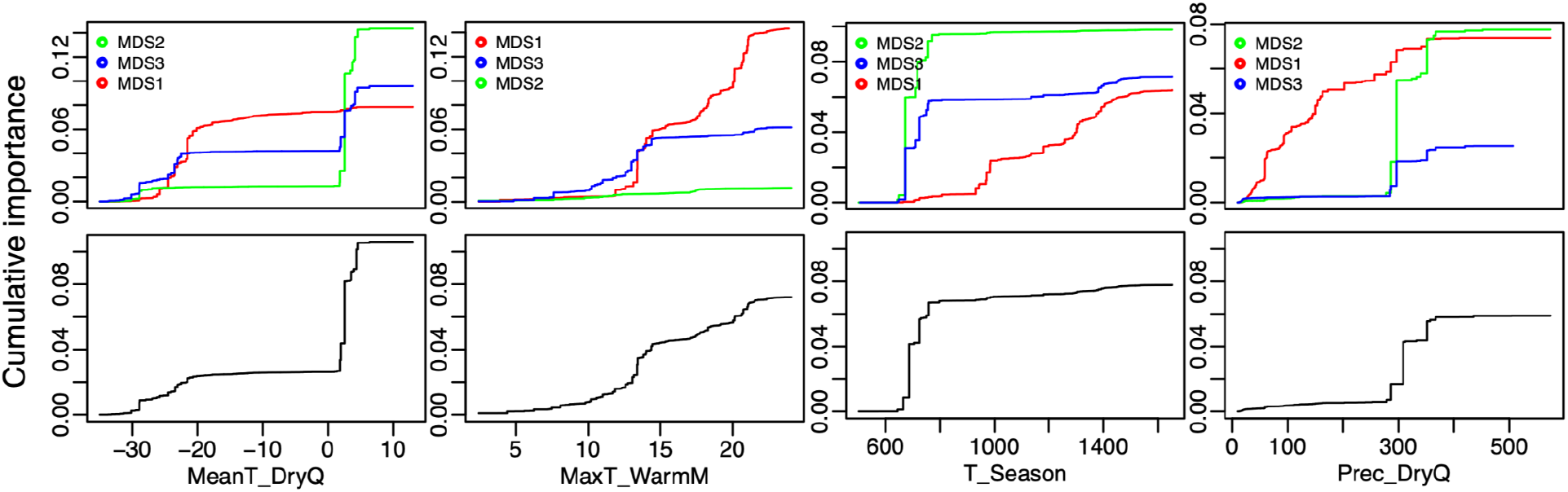
Turnover curves for the three leading gPCs (labeled MDS1, MDS2, MDS3 here) of the wolf dataset. The colored plots in the top row are turnovers of individual gPCs, and the black-and-white plots in the bottom row are their averages. These plots visualize three aspects of the model structure. (1) Which predictor is associated with which gPC. For example, MeanT_DryQ and T_Season are most strongly associated with MDS2 (green line in top-row plots). (2) Where in the predictor range major gPC transitions happen. For example, for MeanT_DryQ the largest transition is just above the freezing point, zero degrees C; which means that wolves’ genetic makeup changes sharply depending on whether it freezes during the driest quarter of the year. (3) Scale of vertical axes shows proportion of variance explained by each predictor for each gPC (top row), and average across all gPCs (bottom row).

Next, we run the *mtry*-based variable selection procedure, relaxing the default passing criteria a bit: we required that a predictor should increase in importance in more that a quarter of all replicates (the default is more than one-third). We relaxed the criteria because our top priority in the end is to obtain the most accurate predictions of adaptation across the landscape. If our main goal was to identify the most important predictors and measure their importance, we would have used more stringent passing criteria (see Discussion). Only three variables did not pass our lenient criteria (Fig. 3 E).

Next, we run 50 *ordinationJackknife* replicates to measure the importance of passing predictors (Fig. 3 F) and to form robust adaptation predictions (averaged across jackknifing replicates) across the landscape. Just like in the simulations above, we formed two types of predictions: direct gPC scores predicted by *randomForest* (“RF predictions”) and turnover-based predictions based on *gradientForest*.

The next step is plotting adaptive neighborhoods and exploring which environmental variables they correspond to (Fig. 5). We begin with exploring unclustered gPC scores predicted by *randomForest* (“RF predictions”). First we plot the PCA of these scores, with points colored according to the values of three PCs (Ellis, Smith, and Pitcher 2012), and project the four selected environmental predictors onto it (Fig. 5 A). We see that blue and green points (landscape locations) are predicted to host wolves adapted to colder and drier conditions compared to warm-colored points, and that wolves at blue points are likely adapted to the high seasonality. The reddest points are associated with the wettest conditions during the dry quarter. We then plotted these predicted points on the landscape (they merge into a continuous layer) and overlaid the points showing the original 94 wolves, colored according to their assigned ecotype (Fig. 5 B, C). We see that distinct colors of the RF predictions tend to align with different ecotypes, with the possible exception of West forest and Atlantic forest ecotypes that (as our model infers) appear to be adapted to similar conditions.

**Figure 5.**
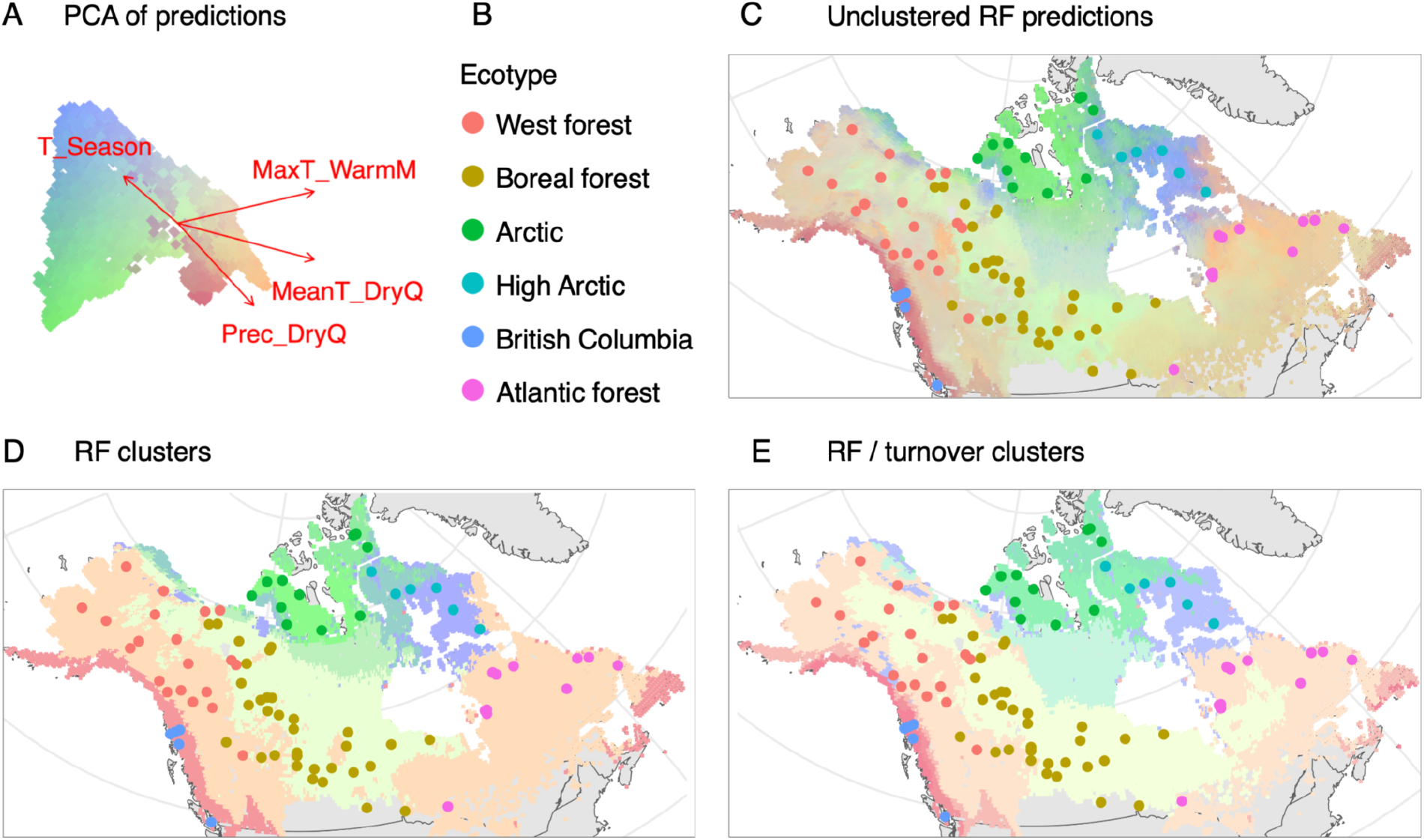
Plotting predicted patterns of differential adaptation for North American wolves. A: PCA of *randomForest* predictions across the landscape, each point being a location within a 37-km-resolution raster covering the region of interest. The four most-important predictors are projected onto the PCA using the function *vegan::envfit*. B: Legend to ecotypes on subsequent panels. C: Unclustered randomForest (RF) predictions, overlaid with the 94 points corresponding to the wolves used for model building (colors identify ecotypes according to panel B). Similarity of colors on the underlying map implies similarity of inferred adaptation. D: Inferring boundaries between adaptive neighborhoods by clustering RF predictions. E: Inferring boundaries between adaptive neighborhoods by clustering turnover curve predictions followed by merging the resulting clusters based on RF predictions.

We next explore the two different options for delineating the boundaries between adaptive neighborhoods. The first one is to cluster the RF predictions directly, which corresponds to clustering of simulation results shown on panels I and K of Fig. 1 and 2. We form 12 clusters and then merge them to get seven or eight. The results look reasonable, cleanly delimiting all the ecotypes with the exception of the High Arctic one, which remains spread across two closely related clusters (Fig. 5 D). Lastly, we do clustering based on turnover curves followed by merging based on RF predictions, like on panels I, L, and M of Fig 1 and 2. Visually, this clustering solution is almost indistinguishable from direct clustering based on RF predictions (Fig. 5 E). Notably, the High Arctic ecotype remains problematic, only this time we see that half of its representatives are attributed to the Arctic cluster.

### Conservation genetics applications

Here we continue with the wolves’ example to demonstrate three practical uses of our methodology: assessing the distribution of risk from climate change across the landscape, planning assisted gene flow, and finding suitable habitat for a specific individual based on its genotype.

The risk of future maladaptation due to upcoming changes in the environment is called a “genetic offset” (Fitzpatrick and Keller 2015). In our method, the genetic offset is simply the Euclidean distance between gPC vectors predicted for the present-day adaptation map (such as on Fig. 5 C) and gPCs predicted using future values of the same predictors. We emphasize that one must use *randomForest* predictions (which are gPC vectors) rather than turnover-based predictions for this calculation, because in a general case involving non-monotony and/or interactions between environmental variables, turnover-based predictions do not accurately reconstruct the distribution of differentially-adapted genotypes on the landscape (Fig. 1 D, L). To make the genetic offset values more interpretable, we scale them to the 90th percentile of gPC distances observed across the present-day landscape.

In this way, the offset of one means that the future will require almost as much additional adaptation as the maximum disparity currently observed between locations. Figure 5 A shows the genetic offset calculated for wolves, relative to climate predictions for 2040-2060: Arctic populations appear to be at the highest risk, while coastal populations are at the lowest risk.

Assisted gene flow is introduction of individuals into a managed population in hopes of introducing genetic variants to help adaptation to the future climate (Aitken and Whitlock 2013). Such individuals should be obtained from location(s) that are *currently* at the state of adaptation that our managed population will need *in the future*. To find such locations, we use the same idea as for the genetic offset calculation: we predict the future gPC vector for our managed population, and compute Euclidean distance between them and present-day gPC predictions across the landscape (Fig. 6 B).

**Figure 6.**
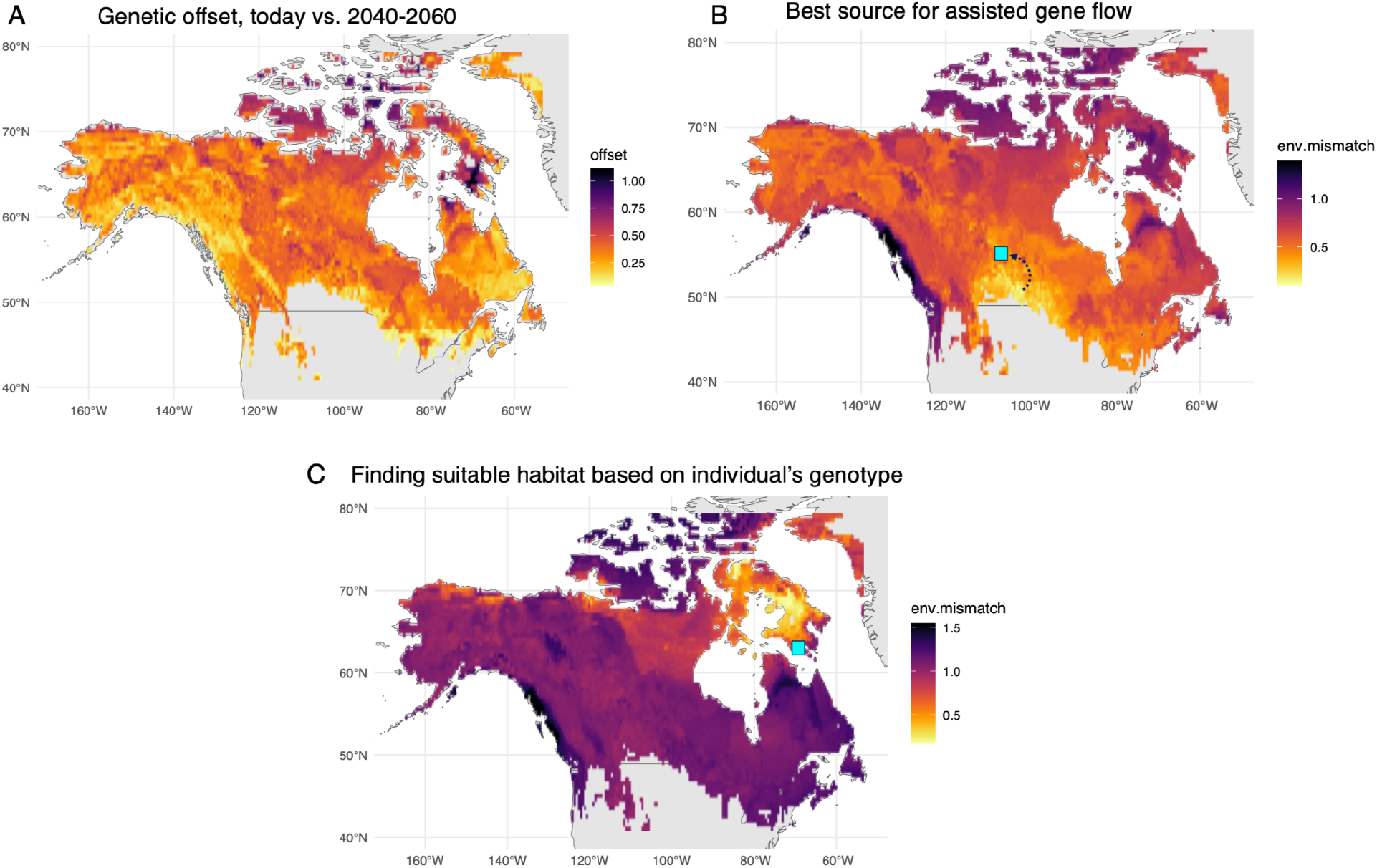
Conservation genetics applications. A: Genetic offset: future maladaptation due to climate change, based on comparison between projected climate (2040-2060) and present day. Arctic is at the highest risk, while both West and East coasts are at the lowest risk. B: Assisted gene flow planning: identify optimal location (origin of the d arrow) to import individuals from to promote adaptation to the future conditions in our focal population (square). C: Finding suitable habitat for a given individual, based on its genotype. Here, one of the wolves (a High Arctic ecotype, sampled at a point indicated by cyan square) was withheld from model-building and then analyzed as an unknown individual to be placed in the most suitable habitat. The model predicted the ecotype-habitat correctly, although the wolf seems to have wandered out of its environmental comfort zone. On all panels, plotted values are scaled to the 90%th percentile of currently observed adaptation disparity across the landscape.

Finding the most suitable location for a given individual (based on its genotype) is a similar problem to the assisted gene flow mapping. We first determine the vector of gPCs for the individual in question by sequencing it and projecting it into the ordination that was used to build the RDAforest model. Once we have the individual’s gPCs, we compute Euclidean distance between them and gPC predictions across the landscape. This mode of analysis can also be used to identify the likely geographic origin of an unknown individual. Our wolf model is quite accurate at placing randomly-withheld individuals into environmental context matching their ecotype (Fig. 6 C).

## Discussion

### Space: the perpetual annoyance

The question how to deal with signals of genetic structure that is exclusively due to spatial separation continues to be a hotly debated issue in the field of genotype-environment association analysis. One such signal is isolation-by-distance (IBD), which is easy to detect by plotting genetic distances against geographic distances (Fig. 3 B). IBD arises through interplay between limited dispersal and local genetic drift, the effects of which cannot be disentangled, i.e., stronger genetic differentiation across a given distance could equally be a result of lower migration and smaller local population sizes (Slatkin 1987; Whitlock and McCauley 1999). To deal with IBD, here we use the fact that it creates the match between the genetic structure as seen in PCA and the geographic map of sampled locations (Novembre et al. 2008; C. Wang, Zöllner, and Rosenberg 2012). One existing method, the “unbundled principal components” (un-PC), already makes use of this fact, detecting putative barriers to migration and long-range admixture events by the distortions they induce in the genetic (PCA-based) map compared to the spatial map (House and Hahn 2018). Conditioning the genetic PCA on spatial coordinates is therefore a valid solution to suppress the IBD signal, assuming there are no physical barriers to migration and/or sizes of local populations do not vary across the landscape. But what these assumptions are violated? The problem is, there is no general way to tell whether such variation is due to physical barriers (which are of no interest to us) or selection against maladapted immigrants (which is what we are looking for): both would generate similar distortions to the genetic structure in PCA space. This means that one should not seek to remove just any signal of spatial patchiness from genetic data. Although it is a relatively common practice to regress any signal of spatial structure that can be captured by Moran eigenvector maps (MEMs) out, we believe that this amounts to splashing the baby out with the bathwater. As we have shown in our simulations (Fig. 1 N and Fig. 2 N), with random forest regression and enough sampling any spatial genetic structure (including instances of true local adaptation) can be captured based on just spatial coordinates. So, if we remove all the detectable spatial signals from the genetic data, we will lose the true adaptation signal as well. The only “space-correction” beyond correction for IBD that we believe is prudent would be for cases when the dataset involves clearly defined physically separated regions, such as different aquifers, mountain tops, or islands, between which migration is likely restricted. These regions should be coded in metadata as dummy quantitative variables (using a column of ones and zeros for each region) and regressed out of the data at the stage of ordination construction (i.e. become additional constraints for the ordination).

### Dealing with collinear predictors

Collinear (correlated) environmental variables are another perpetual problem of ecology. Some prior landscape genomics papers took an ordination-based approach to recode the matrix of correlated environmental variables into uncorrelated environmental PC scores (Bay et al. 2018). This is definitely a valid approach with just one downside: one loses track of the actual variables and so the results become harder to interpret. But should we really strive to minimize correlations among predictors? In classical linear regression the answer is definitely yes, because the regression coefficients for correlated predictors can swing wildly depending on the data and on which predictors are included in the model. In our method this problem is greatly mitigated via extensive resampling of both data and predictors (the staple of the random forest regression) and correlations-aware conditional permutation (Strobl et al. 2008; Ellis, Smith, and Pitcher 2012). Thanks to these features, in our method the inferred predictor importances are stable and the most important predictors are identified correctly, but still their importance remains partially “shared” with correlated predictors. For example, consider the situation when there are two correlated predictors and only one of them is truly influential. This predictor would explain 10% of variation on its own, but including both predictors into the model would spread these 10% among the two depending on the strength of their correlation (maybe 7% for the true predictor and 3% for the correlated one). Whether or not this is a problem depends on the goal of the analysis. If the goal is to identify the drivers of genetic variation across the landscape and measure their importance, then yes, one should try to remove all the correlated but likely non-influential predictors. Conversely, if the goal is to build the most accurate model to predict adaptation across the landscape, retaining correlated predictors will not hurt and may actually help improve the model’s accuracy. The default thresholds in the *mtrySelJack()* function for keeping a predictor are 0.333 for the fraction of replicates in which its importance must increase at higher *mtry*, and 0.1 for the minimum importance relative to the best predictor. More conservative *mtry* selection thresholds (higher importance in 50% or even 75% of high-*mtry* replicates and 0.2 for the minimum allowed importance compared to the best predictor) would be preferred for identifying the most important predictors and measuring the amount of variation they explain. The alternative “lenient mode” for the most accurate model building would be to discard only the most hopeless predictors, those that never increase in importance at higher *mtry* and explain less than 0.01 fraction of variation compared to the best predictor. The possible downside is that the “guided clustering” of neighborhoods based on turnover curves will likely stop working efficiently with too many predictors: a high number of intersecting turnover curves will be splitting the landscape into too many small areas of potential differential adaptation, countering the benefit of integrative nature of each turnover curve. Yet, direct gPC predictions based on *randomForest* are expected to improve, so the lenient mode is definitely worth considering when delineating adaptive neighborhoods. It is advisable to try several levels of predictor selection stringency to see whether relaxing the selection criteria (and retaining more predictors) actually helps improve the overall model accuracy.

### When the true predictor is missing

It is quite possible that the actual driver of adaptation is missing from the variables in *E*, although there might be one or more predictors correlated with it. Since correlated predictors could “stand-in” for the true one, the RDAforest procedure will work, although its predictive power will be limited by the tightness of correlation between true driver and existing predictors. Importantly, there is no formal way to tell whether the most important predictor is indeed the true driver of adaptation or is only correlated with the unknown true driver. Because of this, one should never claim that the true driver has been identified, unless it is validated experimentally, for example in a common garden experiment.

It is also possible to be missing the true driver and not having any predictors correlated with it. In that case, some clearly visible separations of individuals in PCA space would remain unexplained. We seem to have such a situation in the case of Arctic and High Arctic wolves: although they are clearly distinct in the PCA space (Fig. 3 C), our model struggles to predict this (Fig. 5 C-E).

One possible solution to the missing predictor situation, at least as far as building a more accurate predictive model is concerned, is to add spatial coordinates to the table of predictors. If they pass the *mtry*-based selection procedure, that would indicate that they recapitulate the patchiness of genetic structure (Fig. 1 and 2, panel N) better than measured environmental variables, potentially because they are “standing in” for some unknown variable(s). This would be a reasonable solution even after regressing out their linear associations to account for IBD. Still, there is a concern that patches thus detected might be shaped by unrecognized physical barriers to migration rather than by local adaptation. Because of this, such a mode of analysis can only be recommended when there is a good reason to believe that physical connectivity is unrestricted, for example in local-scale seascape genomics.

### Clustering and neighborhood boundaries

Clustering works great to draw neighborhood boundaries in simulations (Fig. 1 and Fig 2), but there is a downside: clustering involves three arbitrary decisions: which clustering method to use (RF-only or turnover+RF), how many clusters to make initially, and which similarity level to merge them at. The function *plot_adaptation()* that does the clustering/merging returns a hierarchical clustering tree with the merging threshold indicated and plots a simple map with resulting clusters numbered on the landscape, both of which can be used as a guide while exploring alternative clustering settings (Fig. 7). However, with no prior data such as ecotype designations that we have for wolves, there is hardly any reliable guidance for choosing clustering settings. We therefore recommend that the practitioners use unclustered maps (such as on Fig. 5 C) as much as possible and avoid making decisions based on the boundaries produced by clustering, unless there is a way to cross-validate the clusters.

**Figure 7.**
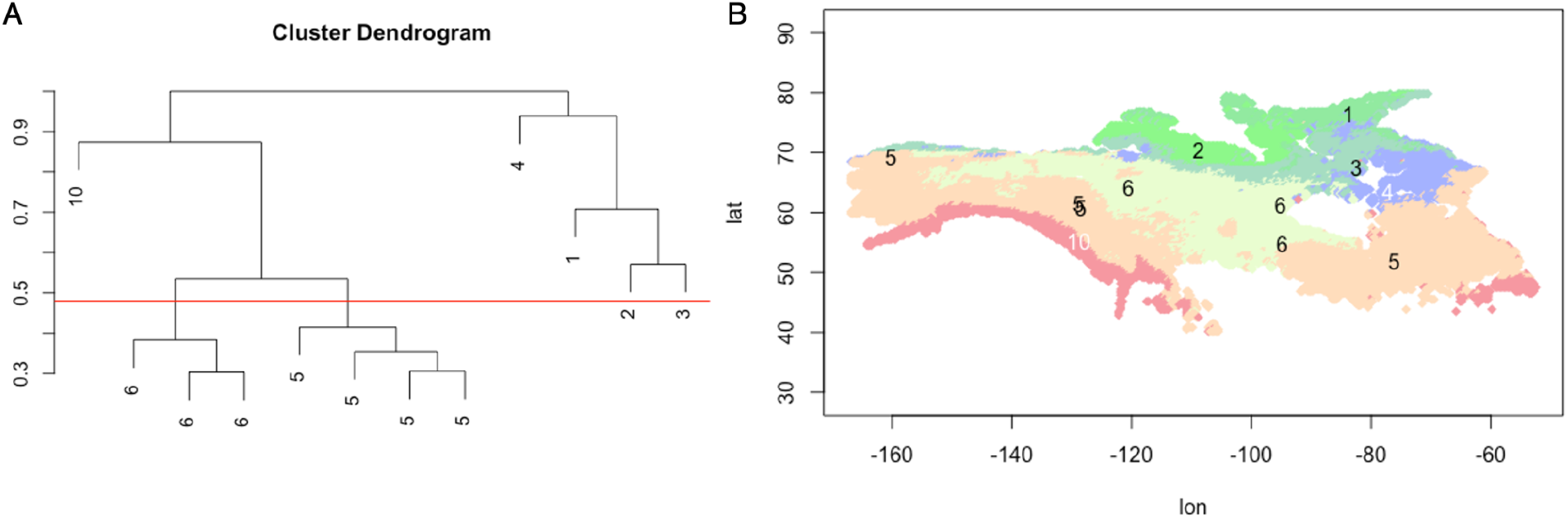
Example clustering-guiding output of the function *plot_adaptation()*. A: Hierarchical clustering tree of RF predictions for cluster medoids. The y-axis indicates how disparate the clusters are; it is scaled in terms of the fraction of maximum observed difference between clusters. The red line is the merging threshold used. Numbers correspond to clusters; merged clusters are assigned the same number. B: Merged clusters on the map. Numbers correspond to clusters on panel A.

### Need for validation

Unfortunately, the results of ecological genomic analyses are rarely validated experimentally, leaving open the question of how reliable are the rankings of putative adaptation drivers and how much can we trust the maps of differential adaptation (such as the ones on Fig. 5). A common garden experiment would test the former: representatives of allegedly differentially adapted populations can be placed under the same condition, where their fitness ranking is expected to reflect the difference between the common-garden environment and the optimal environment predicted for them. Common garden experiments should also be used to validate candidate drivers of adaptation, by measuring fitness of supposedly differentially adapted individuals in two common gardens that are identical except for the focal driver. Maps of adaptive neighborhoods can be validated in a series of reciprocal transplantations between locations that are predicted to require increasingly divergent adaptation. Such experiments are very challenging in animals, but there are few examples of such studies in plants. Two studies, in sorghum (Lasky et al. 2015) and maize (Gates et al. 2019) have successfully validated that environment-associated SNPs do indeed predict how genotypes will fare under specific environmental stressors in a common garden experiment. In poplar, the genetic offset that accounts for unequal importance of environmental variables for adaptation predicted the performance of individuals in a common garden better than “naive” climate difference, which weighs all environmental parameters equally (Fitzpatrick et al. 2021). The most interesting validation thus far has been offered not in plants but in a bird (yellow warbler), where population declines predicted by the future genetic offset corresponded with population declines observed in the field (Bay et al. 2018). This claim was rather extraordinary so it is hardly surprising that it drew criticism (Fitzpatrick, Keller, and Lotterhos 2018). Indeed, genotype-environment association inferences are not always experimentally supported: in European beech, the genomic region supposedly associated with winter temperature adaptation did not show association with fitness in a common garden (Lazic et al. 2024). Overall, the ecological genomics field is still under active development both in terms of experimental design and analytical methods, and experimental validation could greatly help solidify the methodological base, by showing what works and what doesn’t.

## Acknowledgements

This work was supported by the National Science Foundation grant OCE-2318775 to M. V. M. and the NatureNet Science Fellowship by The Nature Conservancy to K. L. B.

## Data and code availability

*RDAforest* method is implemented as a fully documented R package and is available from gitHub at https://github.com/z0on/RDA-forest. The repository also includes installation instructions, summary of main functions, and script and data for the wolf analysis. The output of the wolf Rmarkdown script can be viewed at https://rpubs.com/cmonstr/1268717.

## Author contributions

Both authors contributed ideas that constitute the method. *RDAforest* package was developed by M.V.M, who also wrote the first version of the manuscript. Both authors contributed to finalizing the manuscript.

## References

Aitken, S., and M. Whitlock. 2013. “Assisted Gene Flow to Facilitate Local Adaptation to Climate Change.” Annual Review of Ecology, Evolution, and Systematics 44:367–88.

Balkenhol, Niko, Rachael Y. Dudaniec, Konstantin V. Krutovsky, Jeremy S. Johnson, David M. Cairns, Gernot Segelbacher, Kimberly A. Selkoe, et al. 2017. “Landscape Genomics: Understanding Relationships between Environmental Heterogeneity and Genomic Characteristics of Populations.” In Population Genomics, 261–322. Cham: Springer International Publishing.

Barghi, Neda, Joachim Hermisson, and Christian Schlötterer. 2020. “Polygenic Adaptation: A Unifying Framework to Understand Positive Selection.” Nature Reviews. Genetics 21 (12): 769–81.

Bay, Rachael A., Ryan J. Harrigan, Vinh Le Underwood, H. Lisle Gibbs, Thomas B. Smith, and Kristen Ruegg. 2018. “Genomic Signals of Selection Predict Climate-Driven Population Declines in a Migratory Bird.” Science 359 (6371): 83–86.

Borcard, Daniel, Francois Gillet, and Pierre Legendre. 2011. Numerical Ecology with R. Springer New York.

Capblancq, Thibaut, and Brenna R. Forester. 2021. “Redundancy Analysis: A Swiss Army Knife for Landscape Genomics.” Methods in Ecology and Evolution / British Ecological Society 12 (12): 2298–2309.

Caye, Kevin, Basile Jumentier, Johanna Lepeule, and Olivier François. 2019. “LFMM 2: Fast and Accurate Inference of Gene-Environment Associations in Genome-Wide Studies.” Molecular Biology and Evolution 36 (4): 852–60.

Charlesworth, B., M. Nordborg, and D. Charlesworth. 1997. “The Effects of Local Selection, Balanced Polymorphism and Background Selection on Equilibrium Patterns of Genetic Diversity in Subdivided Populations.” Genetical Research 70 (2): 155–74.

De’Ath, Glenn. 2002. “Multivariate Regression Trees: A New Technique for Modeling Species–environment Relationships.” Ecology 83 (4): 1105–17.

Duruz, Solange, Natalia Sevane, Oliver Selmoni, Elia Vajana, Kevin Leempoel, Sylvie Stucki, Pablo Orozco-terWengel, et al. 2019. “Rapid Identification and Interpretation of Gene-Environment Associations Using the New R.SamBada Landscape Genomics Pipeline.” Molecular Ecology Resources 19 (5): 1355–65.

Ellis, Nick, Stephen J. Smith, and C. Roland Pitcher. 2012. “Gradient Forests: Calculating Importance Gradients on Physical Predictors.” Ecology 93 (1): 156–68.

Fitzpatrick, Matthew C., Vikram E. Chhatre, Raju Y. Soolanayakanahally, and Stephen R. Keller. 2021. “Experimental Support for Genomic Prediction of Climate Maladaptation Using the Machine Learning Approach Gradient Forests.” Molecular Ecology Resources 21 (8): 2749–65.

Fitzpatrick, Matthew C., and Stephen R. Keller. 2015. “Ecological Genomics Meets Community-Level Modelling of Biodiversity: Mapping the Genomic Landscape of Current and Future Environmental Adaptation.” Ecology Letters 18 (1): 1–16.

Fitzpatrick, Matthew C., Stephen R. Keller, and Katie E. Lotterhos. 2018. “Comment on ‘Genomic Signals of Selection Predict Climate-Driven Population Declines in a Migratory Bird.’” Science (New York, N.Y.) 361 (6401). 10.1126/science.aat7279.

Forester, Brenna R., Jesse R. Lasky, Helene H. Wagner, and Dean L. Urban. 2018. “Comparing Methods for Detecting Multilocus Adaptation with Multivariate Genotype--Environment Associations.” Molecular Ecology 27 (9): 2215–33.

Gates, Daniel J., Dan Runcie, Garrett M. Janzen, Alberto Romero Navarro, Martha Willcox, Kai Sonder, Samantha J. Snodgrass, et al. 2019. “Single-Gene Resolution of Locally Adaptive Genetic Variation in Mexican Maize.” bioRxiv. bioRxiv. 10.1101/706739.

House, Geoffrey L., and Matthew W. Hahn. 2018. “Evaluating Methods to Visualize Patterns of Genetic Differentiation on a Landscape.” Molecular Ecology Resources 18 (3): 448–60.

Korneliussen, Thorfinn Sand, Anders Albrechtsen, and Rasmus Nielsen. 2014. “ANGSD: Analysis of Next Generation Sequencing Data.” BMC Bioinformatics 15 (November):356.

Lasky, Jesse R., Hari D. Upadhyaya, Punna Ramu, Santosh Deshpande, C. Tom Hash, Jason Bonnette, Thomas E. Juenger, et al. 2015. “Genome-Environment Associations in Sorghum Landraces Predict Adaptive Traits.” Science Advances 1 (6): e1400218.

Lazic, Desanka, Cornelia Geßner, Katharina J. Liepe, Isabelle Lesur-Kupin, Malte Mader, Céline Blanc-Jolivet, Dušan Gömöry, et al. 2024. “Genomic Variation of European Beech Reveals Signals of Local Adaptation despite High Levels of Phenotypic Plasticity.” Nature Communications 15 (1): 8553.

Legendre, P., and L. Legendre. 2012. Numerical Ecology. Elsevier.

Li, Forrest, Daniel J. Gates, Edward S. Buckler, Matthew B. Hufford, Garrett M. Janzen, Rubén Rellán-Álvarez, Fausto Rodríguez-Zapata, et al. 2024. “The Utility of Environmental Data from Traditional Varieties for Climate-Adaptive Maize Breeding.” bioRxiv. 10.1101/2024.09.19.613351.

Meisner, Jonas, and Anders Albrechtsen. 2018. “Inferring Population Structure and Admixture Proportions in Low-Depth NGS Data.” Genetics 210 (2): 719–31.

Novembre, John, Toby Johnson, Katarzyna Bryc, Zoltán Kutalik, Adam R. Boyko, Adam Auton, Amit Indap, et al. 2008. “Genes Mirror Geography within Europe.” Nature 456 (7218): 98–101.

Oksanen, Jari, Roeland Kindt, Pierre Legendre, Bob O’Hara, M. Henry H. Stevens, Maintainer Jari Oksanen, and Mass Suggests. 2007. “The Vegan Package.” Community Ecology Package 10 (631-637): 719.

Privé, Florian, Keurcien Luu, Bjarni J. Vilhjálmsson, and Michael G. B. Blum. 2020. “Performing Highly Efficient Genome Scans for Local Adaptation with R Package Pcadapt Version 4.” Molecular Biology and Evolution 37 (7): 2153–54.

Schweizer, Rena M., Bridgett M. vonHoldt, Ryan Harrigan, James C. Knowles, Marco Musiani, David Coltman, John Novembre, and Robert K. Wayne. 2016. “Genetic Subdivision and Candidate Genes under Selection in North American Grey Wolves.” Molecular Ecology 25 (1): 380–402.

Sinnott-Armstrong, Nasa, Sahin Naqvi, Manuel Rivas, and Jonathan K. Pritchard. 2021. “GWAS of Three Molecular Traits Highlights Core Genes and Pathways alongside a Highly Polygenic Background.” eLife 10 (February). 10.7554/eLife.58615.

Slatkin, M. 1987. “Gene Flow and the Geographic Structure of Natural Populations.” Science (New York, N.Y.) 236 (4803): 787–92.

Smith, Chris C. R., Gilia Patterson, Peter L. Ralph, and Andrew D. Kern. 2024. “Estimation of Spatial Demographic Maps from Polymorphism Data Using a Neural Network.” Molecular Ecology Resources 24 (7): e14005.

Strobl, Carolin, Anne-Laure Boulesteix, Thomas Kneib, Thomas Augustin, and Achim Zeileis. 2008. “Conditional Variable Importance for Random Forests.” BMC Bioinformatics 9 (July):307.

Wang, Chaolong, Sebastian Zöllner, and Noah A. Rosenberg. 2012. “A Quantitative Comparison of the Similarity between Genes and Geography in Worldwide Human Populations.” PLoS Genetics 8 (8): e1002886.

Wang, Ian J., and Gideon S. Bradburd. 2014. “Isolation by Environment.” Molecular Ecology 23 (23): 5649–62.

Whitlock, M. C., and D. E. McCauley. 1999. “Indirect Measures of Gene Flow and Migration: FST Not Equal to 1/(4Nm + 1).” Heredity 82 (Pt 2) (February):117–25.

Wright, S. 1943. “Isolation by Distance.” Genetics 28 (2): 114–38.

